# TGF-β3 increases the severity of radiation-induced oral mucositis and salivary gland fibrosis in a mouse model

**DOI:** 10.1101/2023.09.12.557315

**Authors:** Ingunn Hanson, Inga Solgård Juvkam, Olga Zlygosteva, Tine Merete Søland, Hilde Kanli Galtung, Eirik Malinen, Nina Frederike Jeppesen Edin

## Abstract

**Purpose:** Toxicities from head and neck (H&N) radiotherapy (RT) may affect patient quality of life and can be dose-limiting. Proteins from the transforming growth factor beta (TGF-β) family are key players in the fibrotic response. While TGF-β1 is known to be pro-fibrotic, TGF-β3 has mainly been considered anti-fibrotic. Moreover, TGF-β3 has been shown to act protective against acute toxicities after radio- and chemotherapy. In the present study, we investigated the effect of TGF-β3 treatment during fractionated H&N RT in a mouse model.

**Materials and methods:** 30 C57BL/6J mice were assigned to three treatment groups. The RT + TGF-β3 group received local fractionated H&N RT with 66 Gy over five days, combined with TGF-β3-injections at 24-hour intervals. Animals in the RT reference group received identical RT without TGF-β3 treatment. The non-irradiated control group was sham-irradiated according to the same RT schedule. In the follow-up period, body weight and symptoms of oral mucositis and lip dermatitis were monitored. Saliva was sampled at five time points. The experiment was terminated 105 days after the first RT fraction. Submandibular and sublingual glands were preserved, sectioned, and stained with Masson’s trichrome to visualize collagen.

**Results:** A subset of mice in the RT + TGF-β3 group displayed increased severity of oral mucositis and increased weight loss, resulting in a significant increase in mortality. Collagen content was significantly increased in the submandibular and sublingual glands for the surviving RT + TGF-β3 mice, compared with non-irradiated controls. In the RT reference group, collagen content was significantly increased in the submandibular gland only. Both RT groups displayed lower saliva production after treatment compared to controls. TGF-β3 treatment did not impact saliva production.

**Conclusions:** When repeatedly administered during fractionated RT at the current dose, TGF-β3 treatment increased acute H&N radiation toxicities and increased mortality. Furthermore, TGF-β3 treatment may increase the severity of radiation-induced salivary gland fibrosis.

## Introduction

Radiotherapy (RT) is an important modality for treatment of head and neck (H&N) cancer, but radiation-induced damage to normal tissue may affect the quality of life of patients and can be dose-limiting. Toxicities from H&N RT may be acute or chronic, and common side effects include dermatitis, oral mucositis, and replacement of salivary gland tissue by connective tissue followed by salivary gland dysfunction and hyposalivation (Sroussi et al. 2017; Siddiqui and Movsas 2017; Wang and Tepper 2021).

The transforming growth factor beta (TGF-β) family of proteins is a group of ubiquitously expressed, pleiotropic proteins with functions within diverse areas including cell cycle regulation, tissue homeostasis, and immunity (Laverty et al. 2009; Fujio et al. 2016; Du et al. 2023). The three isoforms found in humans, TGF-β1-3, possess high structural conformity but exert different functions in several biological processes, including radiation response (Hanson et al. 2023).

TGF-β1 is activated by ionizing radiation and indirectly opposes the effect of cancer radiotherapy through improved DNA repair, suppression of anti-tumor immunity, and remodeling of the tumor microenvironment which can support tumor progression (Barcellos-Hoff 2022). Treatment with recombinant TGF-β3 has been linked to reduced severity of acute radiation damage *in vitro* (Robson et al. 1997) and *in vivo* (Potten et al. 1997; Booth et al. 2000). Furthermore, topical TGF-β3 treatment was developed as a mitigator of acute chemotherapy-induced oral mucositis and showed promise in preclinical trials, but failed phase II clinical trials due to a lack of effectiveness compared to placebo (Sonis et al. 1994; Sonis et al. 1997; Wymenga et al. 1999; Foncuberta et al. 2001).

Radiation-induced fibrosis develops when reactive oxygen species produced by ionizing radiation causes prolonged inflammation, triggering the transformation of local fibroblasts into myofibroblasts and the subsequent excessive deposition of extracellular matrix components (Straub et al. 2015). TGF-β1 is considered a key molecular player in fibrogenesis through fibroblast activation and TGF-β2 is believed to have a similar function (Frangogiannis 2020). Although the role of TGF-β3 in the development of fibrosis is more debated (Wilson 2021), several studies report an anti-fibrotic function (Ask et al. 2008; Bush et al. 2010; Karamichos et al. 2014; Wu et al. 2015). Additionally, treatment with TGF-β3 has been observed to delay the onset and decrease the severity of radiation-induced pulmonary fibrosis in mice (Xu et al. 2014).

Here, we examined if daily TGF-β3 injections could alleviate radiation-induced healthy tissue toxicity during a fractionated RT regime in a mouse model. Surprisingly, we observed increased acute oral toxicity, increased weight loss, and decreased survival in the TGF-β3 treated group. In addition, we found indications of fibrogenesis through increased amount of collagen in the submandibular and sublingual salivary glands in the surviving RT + TGF-β3 treated animals. In the RT reference group, the increase in collagen was significant in the submandibular gland only.

## Materials and Methods

### Animals and treatment

All experiments involving animals were approved by the Norwegian Food Safety Authority according to Directive 2010/63/EU on the protection of animals used for scientific purposes (FOTS ID 27931). Female C57BL/6J mice were purchased from Janvier (France) at nine weeks of age and randomly assigned into three treatment groups: Treatment with TGF-β3 and fractionated RT (RT + TGF-β3) (n=10), fractionated RT only (RT reference) (n=10) and non-irradiated controls (n=10). Data for the non-irradiated controls and the RT reference group have previously been published in (Juvkam et al. 2022) and (Juvkam et al. 2023). A detailed description of the treatment setup is available in (Juvkam et al. 2022). Briefly, animals were anesthetized and placed in a holder with a custom-built lead collimator defining a radiation field covering the oral cavity, pharynx, and major salivary glands. They were then irradiated with 100 kV x-rays at a dose rate of 0.66 Gy/min in 10 fractions of 6.6 Gy over 5 days (total dose 66 Gy). Non-irradiated controls were anesthetized and sham irradiated according to the same schedule. In the follow-up period of 105 days, the body weight and general welfare of the animals were monitored frequently. Every third day during the first 35 days, blinded oral examinations were executed under anesthesia to assess symptoms of acute skin toxicity and oral mucositis, according to criteria developed by the Radiation Therapy Oncology Group (Trotti et al. 2000; Sonis et al. 2004; Mallick et al. 2016). Saliva was sampled on day -8 (baseline), 35, 80, and 105 after the first fraction. Upon indication of moderate to severe oral pain for a subset of animals, all animals were fed DietGel® (ClearH2O, Westbrook, ME, USA), and received analgesic treatment in the form of subcutaneous buprenorphine injections (Temgesic, Indivior, Richmond, VA, USA) three times per day for four consecutive days (day 12 to day 15). Weight loss of more than 20% of baseline weight was defined as a humane endpoint and animals with such weight loss were euthanized by cervical dislocation. On day 105, the remaining animals were euthanized by intraperitoneal (i.p.) injection of anesthetic (Pentobarbital, Exagon® Vet, Richter Pharma AG, Austria) under terminal anesthesia to avoid H&N tissue damage.

### TGF-β3 treatment

Animals in the RT + TGF-β3 group received five i.p. injections of 0.1 ml recombinant TGF-β3 (R&D Systems, MN, USA) at a concentration of 2.5 μg/mL, resulting in a dose of approximately 12.5 μg/kg body weight. Injections were given at 24-hour intervals, with the first administration 24 hours before the first radiation fraction.

### Histological evaluations

Submandibular and sublingual salivary glands were collected immediately after euthanasia. The tissues were then formalin fixed, dehydrated, paraffin-embedded, and sectioned as described in (Juvkam et al. 2022), before staining with Masson’s Trichrome Stain Kit (Abcam, UK) according to the manufacturer’s instructions to identify collagen. Sections were examined using a Nikon E90i microscope and images were acquired with a Nikon DS-Ri1 camera and a CFI Plan Fluor x10 (NA 0.30) objective (Nikon, Japan). Images were analyzed using the Color Threshold function in ImageJ (Schneider et al. 2012) to calculate the ratio of collagen to total tissue area. A detailed image analysis protocol can be found in Supplementary File 1.

### Statistical analyses

The survival data in Figure 1 were analyzed using the log-rank test. Correlations in Figures 2F, 4B, and 4C were analyzed using Kendall’s τ test. The collagen content data in Figure 3G – H were analyzed using one-way ANOVA followed by Tukey’s HSD test. The saliva volume data in Figure 4A was analyzed using repeated measures ANOVA. All comparisons used a significance level of 0.05.

**Figure 1.**
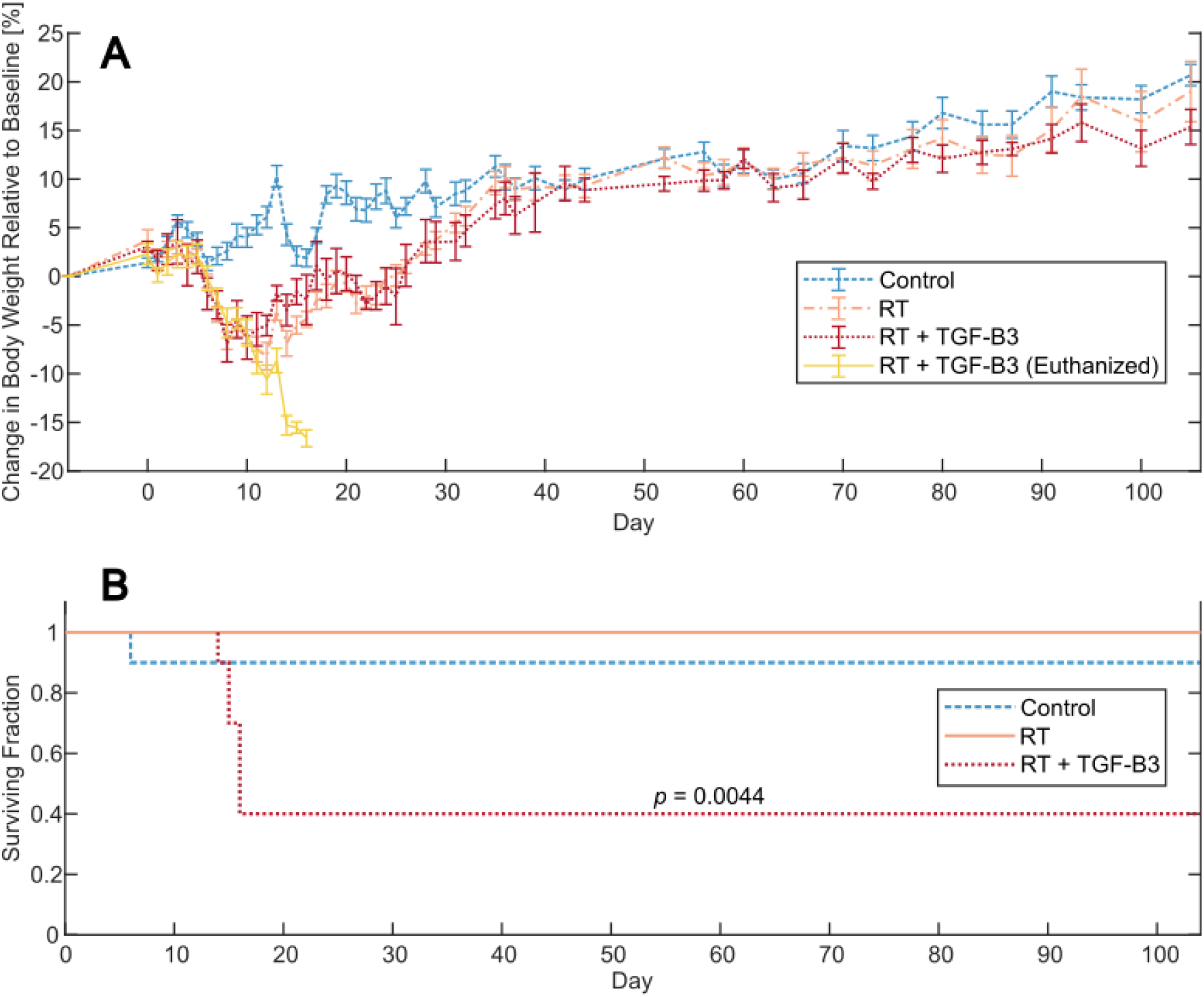
**A**: Change in body weight relative to day 0 for mice in the non-irradiated control (blue), radiotherapy (RT) control (orange), and RT + transforming growth factor beta 3 (TGF-β3) treated groups. A subset of mice in the RT + TGF-β3 group experienced severe weight loss and was euthanized (yellow), while the others recovered (red). **B**: Kaplan Meyer survival plot of the non-irradiated control, RT reference, and RT + TGF-β3 groups. Survival in the RT + TGF-β3 group was significantly lower than the others (p=0.0044, log-rank test).

**Figure 2.**
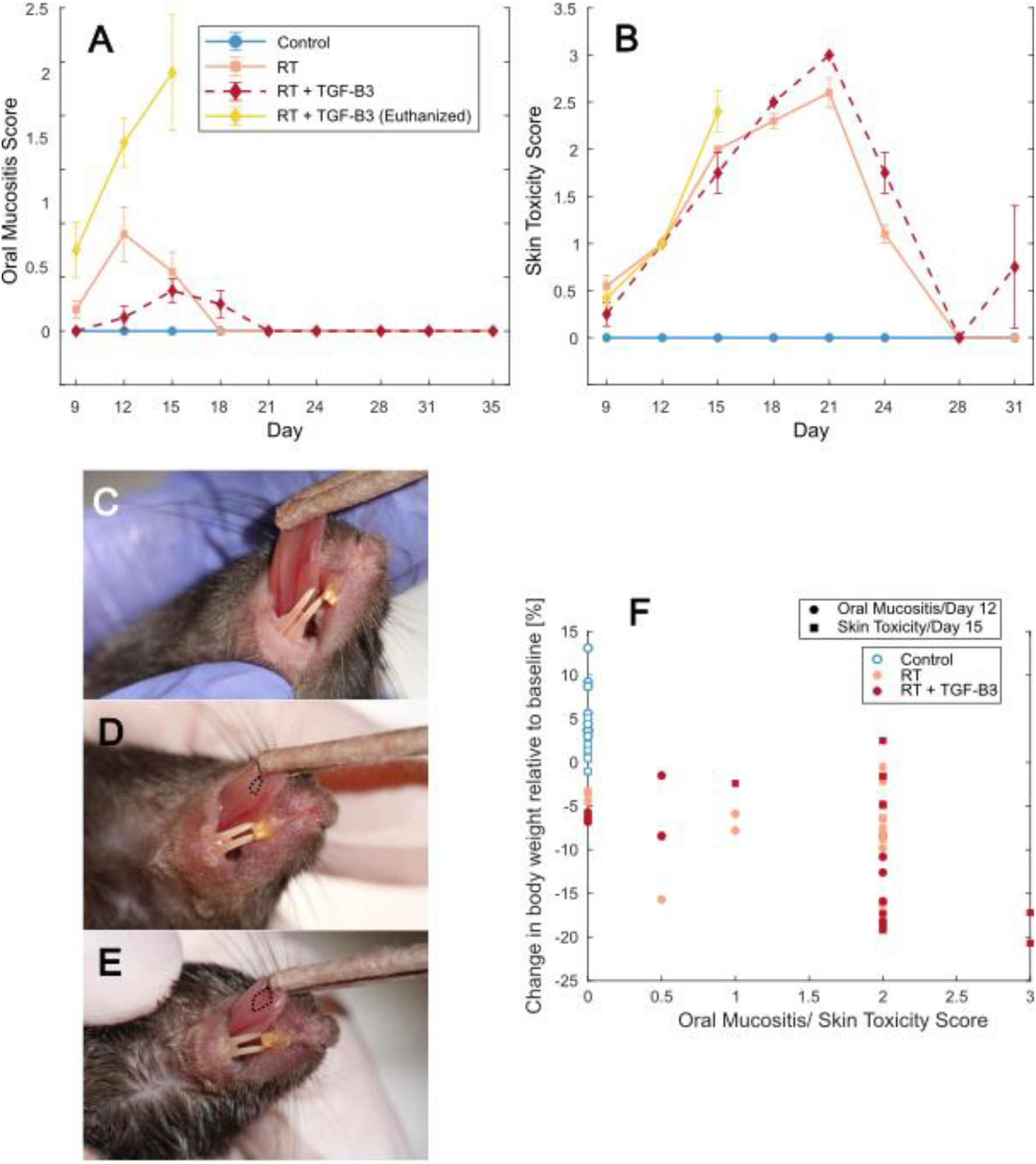
**A:** Oral mucositis score for mice in the non-irradiated control (blue), RT reference (orange), and RT + TGF-β3 groups (euthanized animals in yellow, survivors in red). **B**: Skin toxicity scores for the same groups. **C-E**: Representative images of oral mucositis and skin toxicity in non-irradiated control, RT reference, and RT + TGF-β3 groups, respectively. Outlined area marks oral mucositis. **C**: oral mucositis score 0, skin toxicity score 0. **D**: oral mucositis score 2, skin toxicity score 2. **E**: oral mucositis score 3, skin toxicity score 3. **F**: Correlation between relative change in body weight compared to day 0 and oral mucositis score on day 12 (circles) (p=8.6e-06, Kendall’s τ), and relative change in body weight and skin toxicity score on day 15 (squares) (p=2.8e-05, Kendall’s τ) in non-irradiated controls (blue), RT reference (orange) and RT + TGF-β3 treated mice (red).

**Figure 3.**
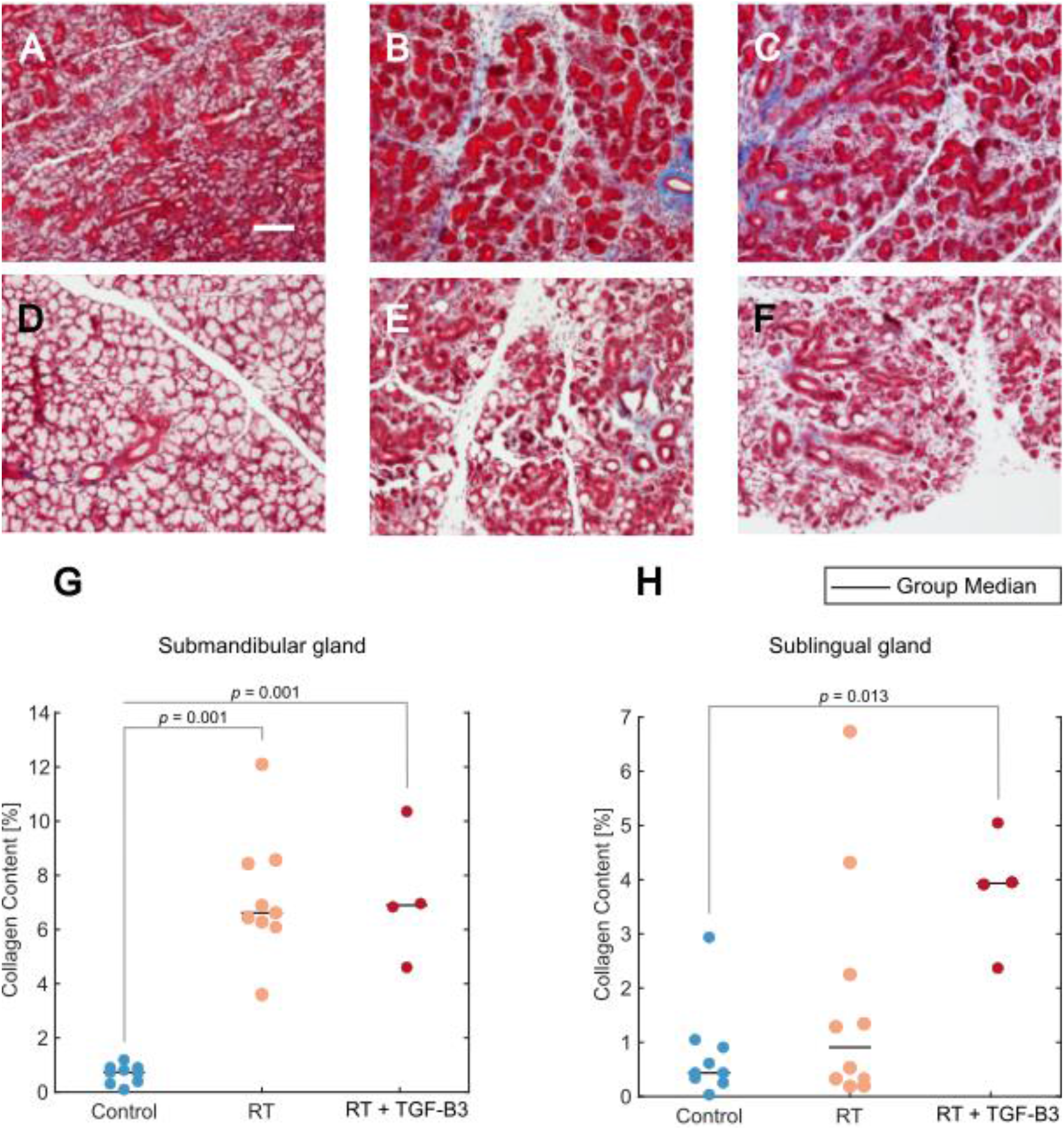
**A-F:** Representative images of tissue sections stained with Masson’s trichrome. Scale bar = 100 μm. **A**: Submandibular salivary gland from non-irradiated control. **B**: Submandibular gland from RT reference group. **C**: Submandibular gland from RT + TGF-β3 treated group. **D**: Sublingual salivary gland from non-irradiated control. **E**: Sublingual gland from RT reference group. **F**: Sublingual gland from RT + TGF-β3 group. **G**: Collagen content in submandibular gland for non-irradiated control (blue), RT reference (orange), and RT + TGF-β3 groups (red) (p=0.001, ANOVA w/Tukey’s HSD). **H**: Collagen content in sublingual gland for non-irradiated control (blue), RT reference (orange), and RT + TGF-β3 groups (red) (p=0.013, ANOVA w/Tukey’s HSD). **G-H**: black line = group median.

**Figure 4.**
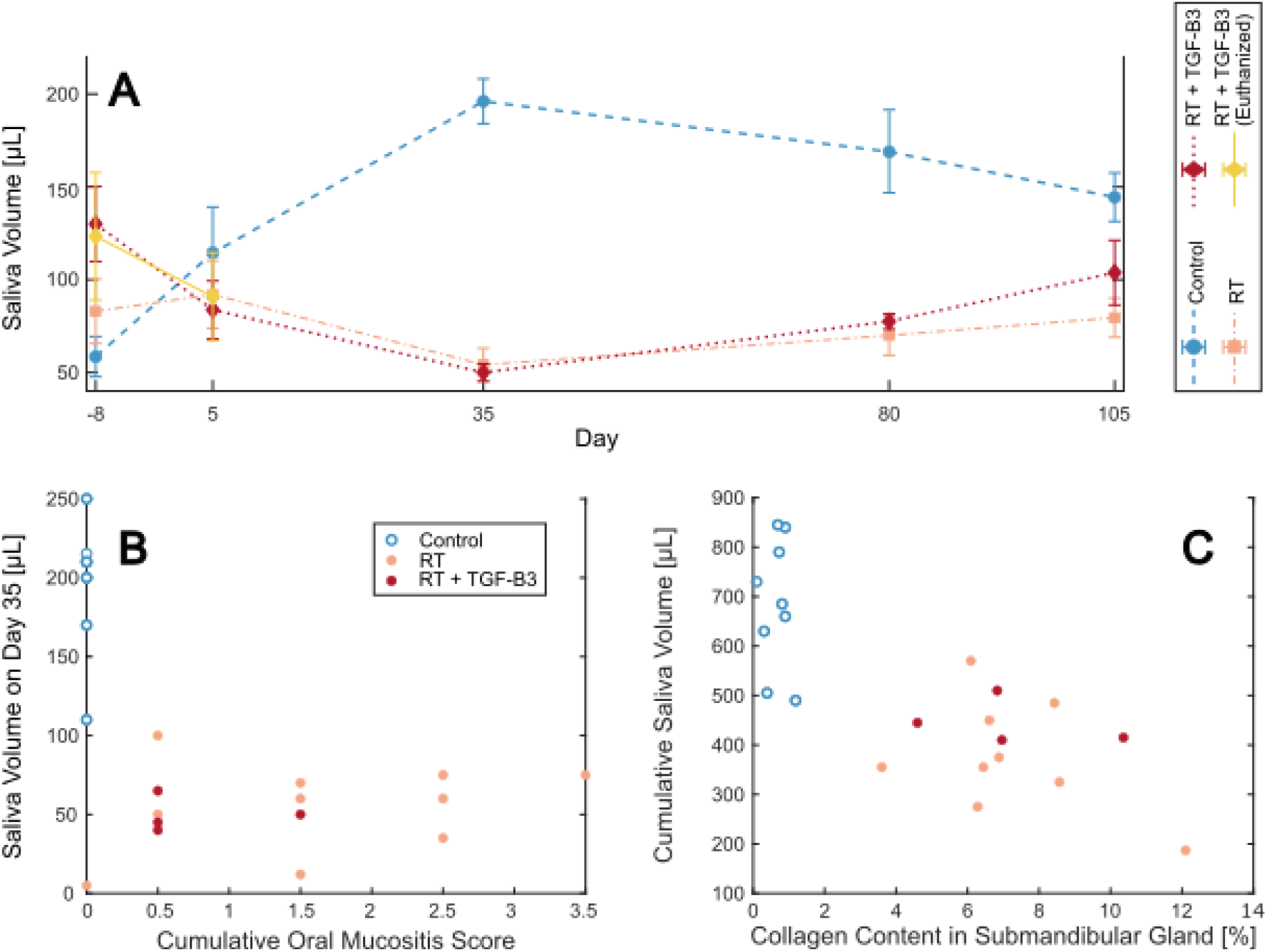
A Saliva volume sampled from mice in the non-irradiated control (blue), RT reference (orange), and RT + TGF-β3 groups (Euthanized mice in yellow, surviving mice in red). Time-dependent increase in non-irradiated controls was significant (p=0.00015, repeated measures ANOVA). B: Saliva volume measured on day 35 correlated with cumulative oral mucositis score up to day 35 (p=0.0092, Kendall’s τ). Non-irradiated controls (blue), RT reference group (orange) and RT + TGF-β3 treated mice (red). C: Cumulative saliva volume up until day 105 correlated with collagen content in the submandibular salivary gland (p=0.00058, Kendall’s τ) for the same groups as B.

## Results

### TGF-β3 treatment increased the mortality after 66 Gy fractionated H&N RT

Body weight was monitored frequently throughout the experiment. Both groups of irradiated mice rapidly lost weight from around day 6, while non-irradiated controls were stable or gaining weight (Figure 1A). Animals in the RT reference group reached a body weight minimum around day 12 before they started gaining weight. A subset of six mice in the RT + TGF-β3 group lost more than 20 % body weight compared to baseline and were euthanized on day 14-16, resulting in a significantly lower survival in this group compared to non-irradiated and RT reference groups (Figure 1B) (p=0.0044, log-rank test). The remaining mice in the RT+TGF-β3 group followed a similar pattern of weight loss as the RT reference group and started gaining weight around day 11. After the acute period, all irradiated animals including the surviving TGF-β3 treated mice followed the same pattern of stable weight gain as the non-irradiated controls.

### TGF-β3 treatment increased the severity of acute oral mucositis

Blinded oral exams were conducted every three days in the acute phase to assess the severity of radiation-induced oral mucositis and skin toxicity. Animals in the RT reference group developed mild to moderate oral mucositis from day 12 (Figure 2A). The RT + TGF-β3 grouped into two subsets: 1) mice with severe oral mucositis and weight loss prior to euthanasia and 2) mild to moderate oral mucositis in the survivors, although their symptoms persevered slightly longer than in the RT reference group (day 21 versus 18). For skin toxicity, animals in the RT reference group and all mice in the RT + TGF-β3 group experienced similar progression (Figure 2B). Skin toxicity was present on day 9 and progressed until day 21 before the symptoms declined and were resolved by day 28 for most individuals. One surviving mouse in the RT + TGF-β3 group experienced severe skin toxicity on the last day of grading (day 31), after exhibiting no signs of skin toxicity on day 28. All non-irradiated controls showed no oral mucositis or skin toxicity. Representative images of mice in the non-irradiated, RT reference, and RT + TGF-β3 groups are displayed in Figure 2C-E.

Relative change in body weight was confirmed to correlate with oral mucositis score on day 12 (p < 0.001, Kendall’s τ) and skin toxicity score on day 15(p < 0.001, Kendall’s τ) (Figure 2F).

### TGF-β3 treatment impacted salivary gland collagen content

Submandibular and sublingual salivary glands were harvested, preserved, and stained with Masson’s trichrome staining after the termination of the experiment on day 105 (Figure 3A-F). Quantitative analysis of submandibular glands revealed increased collagen content in the RT reference and RT + TGF-β3 treated groups, compared to non-irradiated controls (p = 0.001, ANOVA w/ Tukey’s HSD) (Figure 3G). In the sublingual glands, one non-irradiated control mouse was found to have a higher collagen content than the others in the group (Figure 3H). In the RT reference group, three mice displayed increased collagen content, but the increase relative to controls was not significant for the group as a whole. All four RT + TGF-β3 treated survivors displayed increased collagen content compared to non-irradiated controls, and the change relative to controls was significant (p = 0.013, ANOVA w/ Tukey’s HSD). The difference between animals in the RT reference group and the RT + TGF-β3 treated group was not significant.

### TGF-β3 treatment did not impact saliva production

Saliva was collected from anesthetized mice and measured at five time points during the experimental period (Figure 4A). In non-irradiated control mice, saliva production significantly increased during the experiment period (p < 0.001, repeated measures ANOVA). In the RT reference group and surviving RT + TGF-β3 treated mice, there was no increase over time, and saliva volume remained lower than that of control mice until termination. No difference was observed by treatment with TGF-β3 compared to animals in the RT reference group.

Saliva volume measured at the end of the acute period after irradiation (day 35) was significantly negatively correlated to the cumulative oral mucositis score each mouse obtained during the same period (p < 0.001, Kendall’s τ) (Figure 4B).

When the saliva volume measured throughout the experiment was pooled for each animal, this cumulative volume was significantly negatively correlated to the collagen content in the submandibular gland (p < 0.001, Kendall’s τ) (Figure 4C). A similar trend was observed for cumulative saliva volume and collagen content in the sublingual gland, but the correlation was not significant (p=0.48, Kendall’s τ) (Supplementary File 2).

## Discussion

The purpose of the current study was to evaluate the protective effects of TGF-β3 against acute and late radiation toxicities following fractionated radiotherapy in a mouse model. Surprisingly, TGF-β3 treated animals displayed increased oral toxicity, increased weight loss, and decreased survival. Furthermore, TGF-β3 treated animals showed indications of increased fibrogenesis in the sublingual salivary gland.

Repeated injections of TGF-β3 concurrently with fractionated RT increased oral mucositis scores and increased weight loss in a subset of animals, resulting in increased mortality compared to RT alone. Weight loss above 20% of baseline was used as a humane endpoint, thereby influencing the mortality measured in each group. Before euthanasia, administration of analgesic injections and gel fodder were attempted as alleviating measures to avoid unnecessary mortality. When data from all animals were compared, the relative change in body weight was significantly correlated with oral mucositis score on day 12 and skin toxicity score on day 15. From this, we conclude that the severe weight loss observed was a result of pain experienced due to acute radiation toxicity leading to a reduction in food consumption and that severe weight loss was a reasonable humane endpoint for the current study.

In contrast to the results presented here, treatment with TGF-β3 has previously been found to decrease acute radiation toxicity in mouse intestines (Potten et al. 1997; Booth et al. 2000). Furthermore, topical application of TGF-β3 in a Syrian golden hamster model decreased the severity of chemotherapy-induced oral mucositis through inhibition of epithelial cell proliferation, thereby increasing survival (Sonis et al. 1994; Sonis et al. 1997). Mouthwash with TGF-β3 as the active ingredient was tested in cancer patients receiving chemotherapy, but failed to document an effect compared to placebo and did not progress beyond phase II clinical trials (Wymenga et al. 1999; Foncuberta et al. 2001). However, no clinically relevant adverse effects of TGF-β3 treatments were observed in any of these studies.

Alleviation of acute side effects by TGF-β3 in previous studies was attributed to the induction of a transient cell cycle arrest by inhibition of G1/S progression, allowing stem cells in regenerating tissues time for DNA damage repair and thereby reduce radiation damage (Khalil et al. 1994; McCormack et al. 1997; Robson et al. 1997; Nakamura et al. 2010). In the studies referred to above, however, a protective effect was seen when TGF-β3 treatment was administered before challenge with ionizing radiation or chemotherapy. Potten et al. observed that compared to a treatment regime with five i.p. TGF-β3 injections before irradiation, the addition of one post-irradiation injection decreased the radio-protective effect on the intestinal crypts from a factor of 4.9 to 1.6 (Potten et al. 1997). Furthermore, when TGF-β3 injections were administered solely post-irradiation, this induced a radio-sensitization and surviving crypts decreased to 1/3 of control. From this, the authors concluded that when the TGF-β3-induced cell cycle arrest was prolonged after irradiation, tissue regeneration was impeded. The increase in acute radiotoxicity from TGF-β3 treatment observed in the current study may result from the administration of TGF-β3 between RT fractions, which may have inhibited the healing process of radiation-damaged oral mucosa through a sustained G1/S cell cycle arrest of stem cells or transit amplifying cells (Cancedda and Mastrogiacomo 2023).

Treatment with RT and TGF-β3 significantly increased collagen content in the submandibular and sublingual salivary glands, whereas RT alone significantly increased collagen content in the submandibular gland only. In the RT group, collagen content in the sublingual gland varied markedly between animals, and the difference between the RT + TGF-β3 group and the RT reference group was not significant for either gland. This suggests that treatment with TGF-β3 may have enhanced a trend that appeared after RT. While TGF-β1 is widely accepted as a fibrogenic growth factor, TGF-β3 has been observed to exert anti-fibrotic functions in vitro and in vivo (Shah et al. 1995; Chang et al. 2014; Karamichos et al. 2014; Wu et al. 2015; Xue et al. 2019; Escasany et al. 2021). In agreement with our results, evidence suggests the role of TGF-β3 as a purely anti-fibrotic cytokine is too simplistic (Wilson 2021; Hanson et al. 2023).

Xu et al. successfully reduced and delayed the development of radiation-induced pulmonary fibrosis in mice through i.p. TGF-β3 injections (Xu et al. 2014). However, the RT and TGF-β3 treatment regimens employed differed from the ones in the present study. Notably, TGF-β3 injections were administered weekly after irradiation until study termination, whereas TGF-β3 treatment in the current study was terminated after the conclusion of the RT. It is possible that to delay and the reduce development of fibrosis, treatment must be continued until or beyond the appearance of pathological changes. Moreover, the dose of TGF-β3 was lower at 1 μg/kg, compared to ∼12.5 μg/kg in the current study. In previous preclinical and clinical studies where TGF-β3 has been demonstrated to reduce scar tissue formation, the treatment has typically been injected locally, and the dose is not directly comparable to the current study (Shah et al. 1995; Lanning et al. 2000; Loewen et al. 2001; Hosokawa et al. 2003; Ferguson et al. 2009; Bush et al. 2010; So et al. 2011; Chang et al. 2014).

Several studies have observed a shift in the balance between endogenously expressed TGF-β1 and TGF-β3 after irradiation and have speculated that this occurrence rather than the absolute concentration of either influences the development of radiation fibrosis (Wang and Robbins 1996; Zhao et al. 2000; Seong et al. 2000). Furthermore, the TGF-β1/TGF-β3 balance has been observed to differ between oral mucosa and cutaneous tissue, possibly explaining the improved healing abilities of the former (Schrementi et al. 2008; Jiajia Zhao 2015). Considering this, the endogenous TGF-β1 concentration and its change after RT, in addition to the administered TGF-β3 dose, may have influenced the development of salivary gland fibrosis in the current model.

TGF-β signaling is complicated by protein latency and receptor binding. To bind to their receptor complexes, TGF-β proteins must first be activated through disassociation from the latency-associated protein. Activation mechanisms include ionizing radiation and biological molecules, including integrins and proteases, and differ between isoforms (Annes et al. 2003). Disparities in the activation states of TGF-β1 and TGF-β3 may influence fibrogenesis as well as other functional effects. Furthermore, both isoforms have been shown to bind to and signal through TGF-β type I receptors activin-like kinase (ALK) 1 and ALK5, with downstream phosphorylation of Smad 1/5/8 and Smad 2/3, respectively, resulting in different functional effects (Lux et al. 1999; Shi and Massagué 2003; De Kroon et al. 2015). In a recent study, we discovered that ALK5 inhibition induced ALK1 binding by TGF-β3, indicating a competition between the two receptors, where ALK5 has the higher affinity (Hanson et al. 2022). Thus the relative availability of different TGF-β receptors, in addition to the concentration of the cytokines, may influence the development of fibrosis mediated by TGF-β.

In non-irradiated control animals, saliva production increased significantly throughout the experiment period. This is in agreement with previous reports in C57BL/6 mice from 10 to 30 weeks of age (Choi et al. 2013). Animals in the RT reference and RT + TGF-β3 groups did not display this increase. Furthermore, saliva volume sampled at the end of the acute period was significantly negatively correlated to acute radiation damage through oral mucositis score, and the total saliva volume measured throughout the experiment for each animal was significantly negatively correlated to collagen content in the submandibular gland. Although these two parameters were not significantly correlated for the sublingual gland, a similar trend was noted here. Thus, the relative decrease in saliva production in irradiated compared to non-irradiated animals was probably a symptom of salivary gland radiation damage and was not affected by treatment with TGF-β3.

To summarize, we hypothesize that the increase in acute toxicity in the form of increased severity of oral mucositis was due to a TGF-β3-induced G1/S cell cycle arrest, which was sustained when TGF-β3 injections were repeated between irradiation fractions, disallowing stem cells or transit amplifying cells to repopulate damaged oral mucosa. The increased sublingual gland collagen content in the TGF-β3 treated animals was possibly influenced by TGF-β3 dose, the ratio of TGF-β3 to endogenous TGF-β1 concentration, or endogenous availability of TGF-β receptors. At the current dose, and with the current treatment regimen, injections with recombinant TGF-β3 did not mitigate fibrosis as expected but increased both acute and late toxicities of H&N RT in C57BL/6J mice.

## Acknowledgements

We thank Delmon Arous for help with the dosimetry measurements.

This work was supported by UiO Life Science at the University of Oslo under grant reference 2018/10221 and South-Eastern Norway Regional Health Authority under grant number 2019050.

