## Supplementary File 1 for "TGF-β3 increases the severity of radiation-induced oral mucositis and salivary gland fibrosis in a mouse model"

**Calculation of collagen fiber area in Masson’s Trichrome stained tissue sections using ImageJ**

1. Remove white background:
   - Image\Adjust\Color Threshold:

Hue: 0-255

Saturation: 0-255

Brightness: 235-255

- Select
- Edit\Clear
- Analyze\Measure (*Total Tissue Area*)

2. Quantify blue pixels:
   - Image\Adjust\Color Threshold:

Hue: 0-255

Saturation: 0-255

Brightness: 235-255

- Select
- Analyze\Measure (*Collagen Area*)

3. $Collagen Ratio=\frac{Collagen Area}{Total Tissue Area}$
