## Supplementary figures and images for "TGF-β3 increases the severity of radiation-induced oral mucositis and salivary gland fibrosis in a mouse model"

### Supplementary File 2

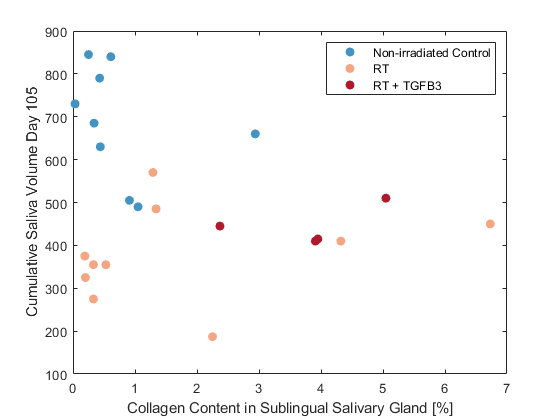
